# Endosomal trafficking protein TBC-2 is required for the longevity of long-lived mitochondrial mutants

**DOI:** 10.1101/2023.02.10.528031

**Authors:** Annika Traa, Hazel Shields, Abdelrahman AlOkda, Zenith D. Rudich, Bokang Ko, Jeremy M. Van Raamsdonk

## Abstract

Mutations that result in a mild impairment of mitochondrial function can extend longevity. Previous studies have shown that the increase in lifespan is dependent on stress responsive transcription factors, including DAF-16/FOXO, which exhibits increased nuclear localization in long-lived mitochondrial mutants. We recently found that the localization of DAF-16 within the cell is dependent on the endosomal trafficking protein TBC-2. Based on the important role of DAF-16 in both longevity and resistance to stress, we examined the effect of disrupting *tbc-2* on lifespan and stress resistance in the long-lived mitochondrial mutants *nuo-6* and *isp-1* in *C. elegans*. Loss of *tbc-2* markedly reduced the long lifespans of both mitochondrial mutants. Disruption of *tbc-2* also decreased resistance to specific exogenous stressors in *nuo-6* and *isp-1* mutants. In contrast, *tbc-2* inhibition had no effect on oxidative stress resistance or lifespan in *isp-1* worms when DAF-16 is absent suggesting that the effect of *tbc-2* on mitochondrial mutant lifespan may be mediated by mislocalization of DAF-16. However, this result is complicated by the fact that deletion of *daf-16* markedly decreases both phenotypes in *isp-1* worms. Surprisingly, disruption of *tbc-2* did not prevent the upregulation of DAF-16 target genes in the long-lived mitochondrial mutants, suggesting the possibility that the effect of *tbc-2* on lifespan and stress resistance in the long-lived mitochondrial mutants is at least partially independent of its effects on DAF-16 localization. Overall, this work demonstrates the importance of endosomal trafficking for the extended longevity and enhanced stress resistance resulting from mild impairment of mitochondrial function.

## Introduction

Mitochondria are membrane-bound organelles that account for most of the energy production in the cell and have important roles in intracellular signaling and metabolism. Despite the importance of mitochondria to cellular function, mild impairment of mitochondrial function can extend lifespan and increase resistance to stress (Dues et al., 2017; Feng et al., 2001; Lakowski and Hekimi, 1996; Schaar et al., 2015; Wong et al., 1995; Wu et al., 2018; Yang and Hekimi, 2010). The ability of mutations affecting the mitochondria to extend lifespan has been demonstrated in multiple species including worms (Feng et al., 2001; Lakowski and Hekimi, 1996; Wong et al., 1995; Yang and Hekimi, 2010), flies (Copeland et al., 2009) and mice (Dell’agnello et al., 2007; Liu et al., 2005), thereby indicating conservation across species.

Among the most well studied mitochondrial mutants in *Caenorhabditis elegans* are *nuo-6* and *isp-1*.The *nuo-6* gene encodes a subunit of Complex I of the mitochondrial electron transport chain (Yang and Hekimi, 2010), while the *isp-1* gene encodes a subunit of the Rieske iron sulfur protein in Complex III of the electron transport chain (Feng et al., 2001). Although a number of studies have identified factors that are important for the long lifespan of these mitochondrial mutants (Baruah et al., 2014; Campos et al., 2021; Harris-Gauthier et al., 2022; Hwang et al., 2014; Jung et al., 2021; Lee et al., 2010; Munkacsy et al., 2016; Senchuk et al., 2018; Walter et al., 2011; Wu et al., 2018; Yee et al., 2014), the mechanism by which mild mitochondrial impairment extends longevity remains incompletely understood.

Among the factors that are required for the longevity of the long-lived mitochondrial mutants is the FOXO transcription factor DAF-16 (Senchuk et al., 2018). *nuo-6* and *isp-1* mutants exhibit increased expression of DAF-16 target genes, which is dependent on DAF-16 and the DAF-16 deubiquitylase MATH-33. DAF-16 is part of the insulin/IGF-1 signaling pathway. Under conditions of decreased insulin/IGF-1 signaling or increased stress, DAF-16 translocates from the cytoplasm to the nucleus to modulate gene expression. Disrupting DAF-16 reverts *nuo-6* and *isp-1* lifespan to wild-type (Senchuk et al., 2018).

We recently showed that cytoplasmic DAF-16 can be localized to endosomes (Meras et al., 2022). Disruption of the insulin/IGF-1 signaling pathway decreases endosomal localization of DAF-16 while increasing its localization to the nucleus. Endosomal localization of DAF-16 is also affected by disruption of endosomal trafficking proteins. Specifically, loss of the Rab GTPase activating protein (GAP) TBC-2 increases the localization of DAF-16 to endosomes, while inhibition of the TBC-2 targets RAB-5 and RAB-7 has the opposite effect (Chotard et al., 2010; Law and Rocheleau, 2017; Meras et al., 2022). The increased endosomal localization of DAF-16 in *tbc-2* mutants results in a decreased ability of DAF-16 to translocate to the nucleus in response to specific stresses (Traa et al., 2022).

Under normal conditions, internalized vesicles containing cargo from the plasma membrane, such as ligand bound receptors, fuse together to form early endosomes, which contain RAB-5. From the early endosome, the internalized cargo can be sorted to the recycling endosome, the Trans-Golgi network or to late endosomes, which contain RAB-7. Cargo transported to late endosomes can then be degraded by the lysosome. When *tbc-2* is disrupted, there is an accumulation of enlarged late endosomes resulting from increased activation of RAB-5 (Chotard et al., 2010).

In this work, we examined the role of TBC-2 in the extended longevity and enhanced stress resistance of long-lived *C. elegans* mitochondrial mutants. We found that disruption of *tbc-2* decreases lifespan and resistance to specific exogenous stressors in *nuo-6* and *isp-1* worms. Surprisingly, the *tbc-2* deletion failed to prevent upregulation of DAF-16 target genes in the long-lived mitochondrial mutants. Overall, this work demonstrates the importance of TBC-2 in stress resistance and longevity and suggests that *tbc-2* disruption may be affecting these phenotypes in long-lived mitochondrial mutants at least partially through DAF-16-independent pathways.

## Results

### Disruption of TBC-2 decreases the lifespan of long-lived mitochondrial mutants

We previously showed that both *daf-2* mutants and the long-lived mitochondrial mutants *nuo-6* and *isp-1* are dependent on DAF-16 for their enhanced longevity (Dues et al., 2019; Senchuk et al., 2018). We also showed that loss of *tbc-2* decreases the lifespan of long-lived *daf-2* insulin-IGF1 receptor mutants (Meras et al., 2022). Since deletion of *tbc-2* affects the ability of DAF-16 to translocate to the nucleus (Traa et al., 2022), we wondered whether the disruption of *tbc-2* would also decrease the lifespan of the long-lived mitochondrial mutants. Accordingly, we crossed *tbc-2* deletion mutants to *nuo-6* and *isp-1* worms and quantified the resulting effect on lifespan. We found that *nuo-6* and *isp-1* worms both exhibit a marked decrease in lifespan when *tbc-2* is disrupted (**Fig. 1**).

**Figure 1.**
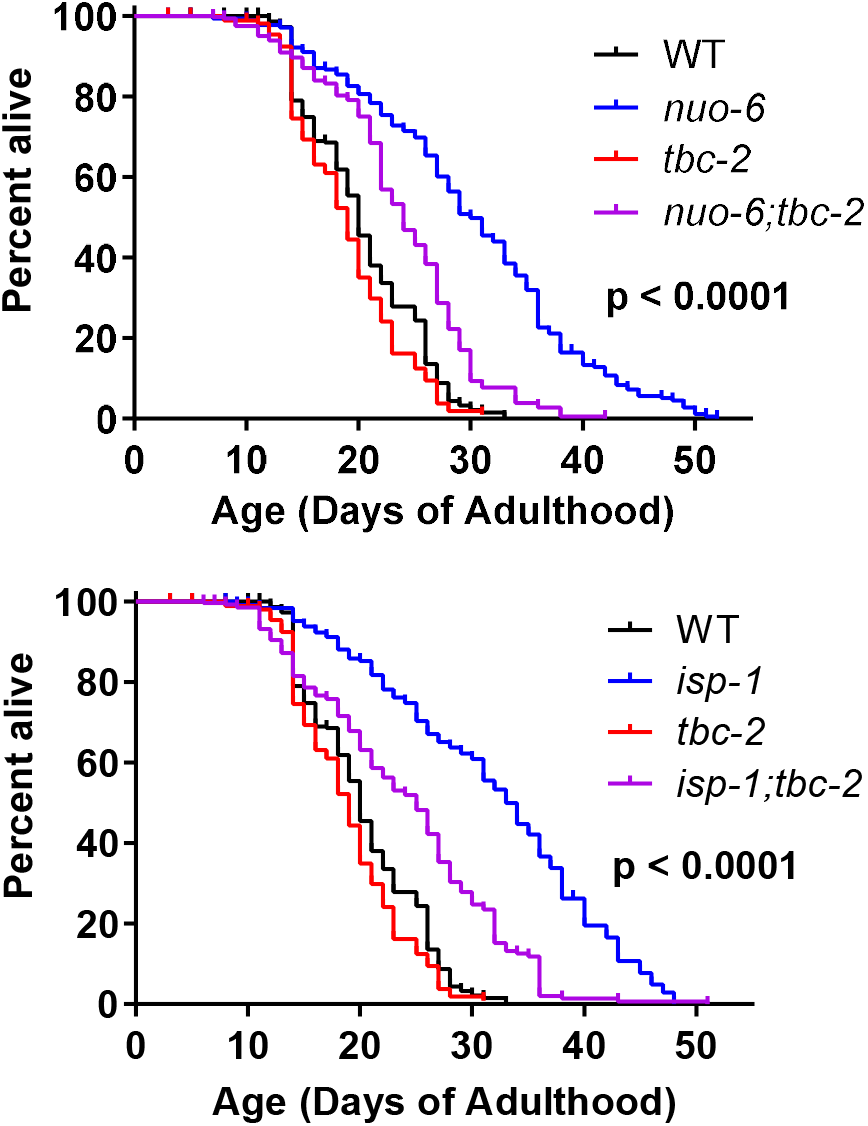
TBC-2 is required for the longevity of long-lived mitochondrial mutants. The long-lived mitochondrial mutants *nuo-6* and *isp-1* were crossed to *tbc-2* deletion mutants. Disruption of *tbc-2* markedly reduced the lifespan of both mutants, while only having a minor impact on wild-type lifespan. Significance indicates differences between blue and purple lifespan curves and was determined using the log-rank test. A minimum of three biological replicates per strain were performed. Raw lifespan data can be found in **Table S1**.

### Disruption of TBC-2 decreases resistance to specific stresses in long-lived mitochondrial mutants

Next, we sought to determine the role of TBC-2 in the enhanced stress resistance of the long-lived mitochondrial mutants and whether this might be contributing to the effect of *tbc-2* disruption on their lifespan. We quantified resistance to chronic oxidative stress (4 mM paraquat), acute oxidative stress (300 μM juglone), heat stress (37°C), bacterial pathogen stress (*P. aeruginosa* strain PA14), osmotic stress (450 mM and 550 mM NaCl) and anoxic stress.

In *nuo-6* mutants, disruption of *tbc-2* significantly decreased resistance to both chronic and acute oxidative stress (**Fig. 2A,B**). In contrast, the loss of *tbc-2* did not significantly decrease resistance to heat stress (**Fig. 2C**), bacterial pathogen stress (**Fig. 2D**), osmotic stress (**Fig. 2E,F**) or anoxic stress (**Fig. 2G**) in *nuo-6* worms. In *isp-1* mutants, deletion of *tbc-2* decreased resistance to chronic oxidative stress (**Fig. 3A**) but did not affect resistance to acute oxidative stress (**Fig. 3B**), heat stress (**Fig. 3C**) or bacterial pathogens (**Fig. 3D**). Disruption of *tbc-2* decreased resistance to osmotic stress in *isp-1* worms (**Fig. 3E,F**), but did not decrease resistance to anoxia (**Fig. 3G**). Combined, these results indicate that disruption of *tbc-2* alters resistance to some, but not all, external stressors in long-lived mitochondrial mutants.

**Figure 2.**
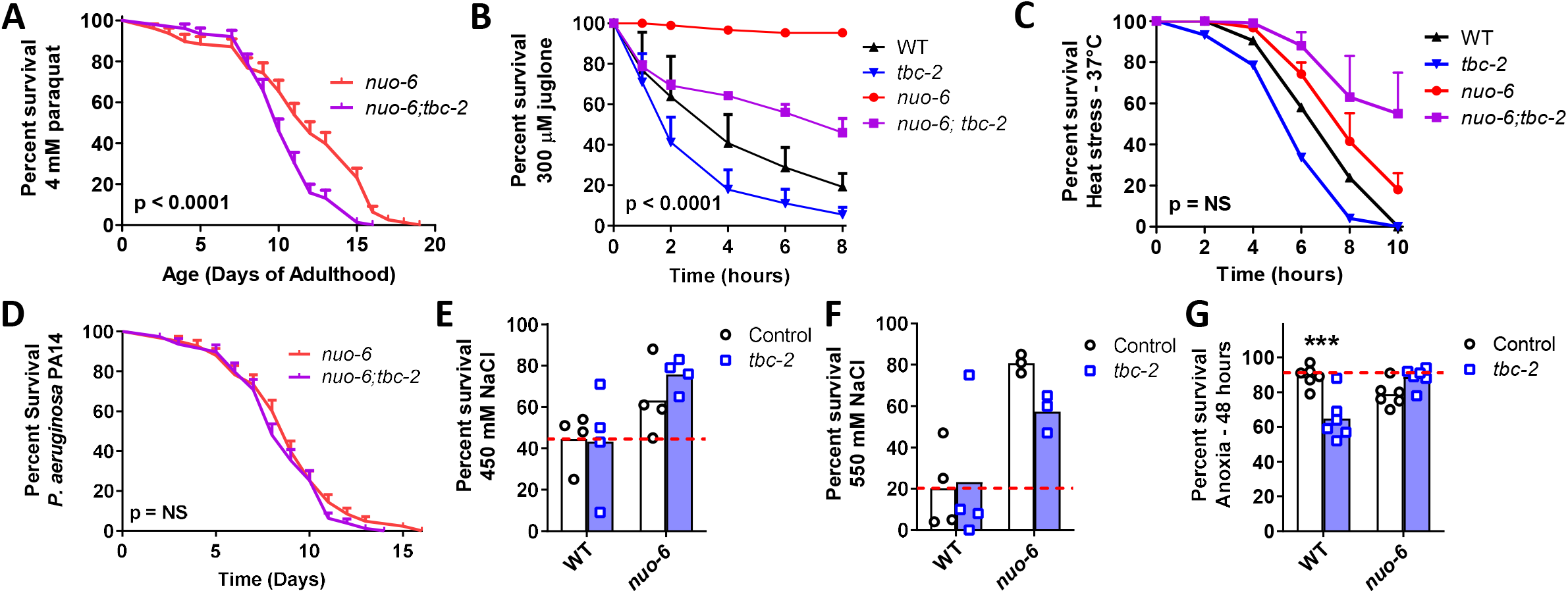
Disruption of *tbc-2* decreases resistance to oxidative stress in long-lived *nuo-6* mutants. Disruption of *tbc-2* decreases resistance to both chronic (**A**; 4 mM paraquat) and acute (**B**; 300 μM juglone) oxidative stress in *nuo-6* worms but has no significant effect on resistance heat stress (**C**; 37°C) in these worms. Disruption of *tbc-2* did not affect resistance to bacterial pathogen stress (**D**; *P. aeruginosa* strain PA14), osmotic stress (**E,F**; 450 mM NaCl, 550 mM NaCl) or anoxia (**G**; 48 hours) in *nuo-6* worms. A minimum of three biological replicates were performed. Statistical significance was assessed using the logrank test in panels A and D, a repeated measures ANOVA in panels B and C, and a two-way ANOVA with Šidák’s multiple comparison test in panels E-G. p values indicate significance of difference between *nuo-6* (red line) and *nuo-6;tbc-2* (purple line). Error bars indicate SEM. ***p<0.001.

**Figure 3.**
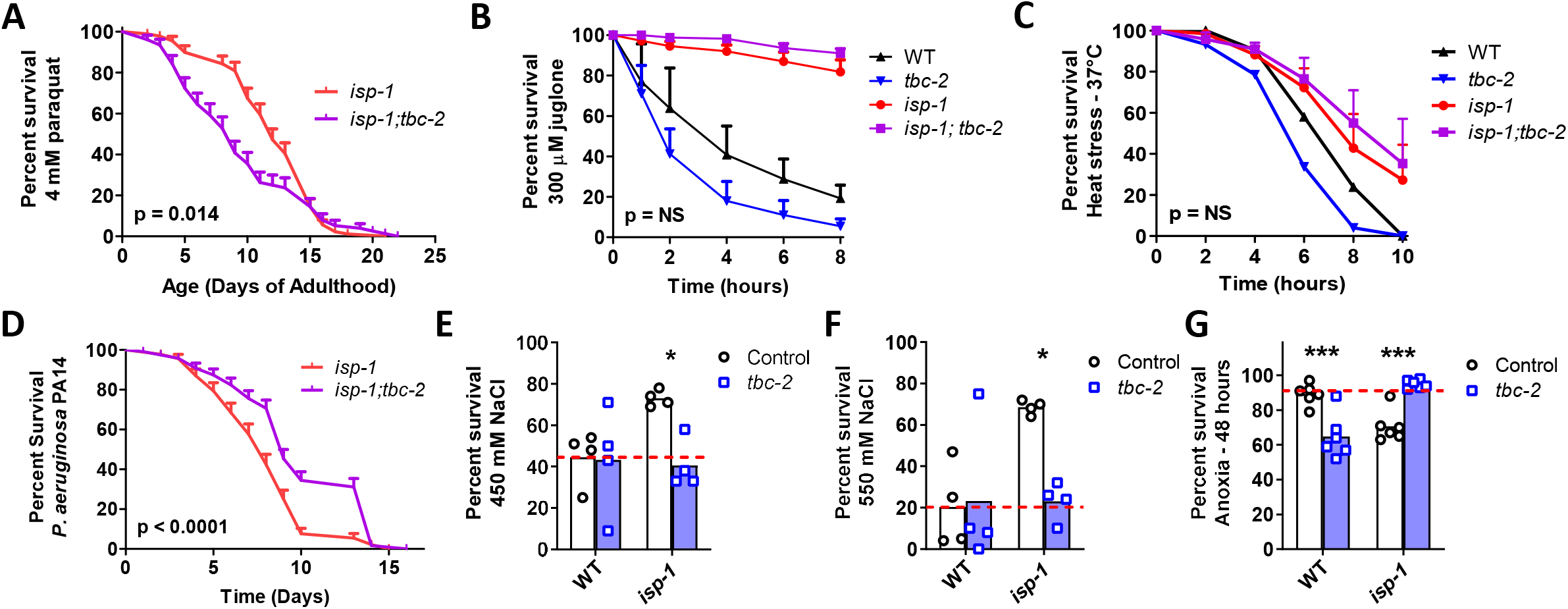
Disruption of *tbc-2* decreases stress resistance in long-lived *isp-1* mutants. Disruption of *tbc-2* decreases resistance to chronic oxidative stress in *isp-1* worms (**A**; 4 mM paraquat) but has no effect on resistance to acute oxidative stress (**B**; 300 μM juglone) or heat stress (**C**; 37°C) in these worms. The *tbc-2* deletion increases resistance to bacterial pathogen stress in *isp-1* worms (**D**; *P. aeruginosa* strain PA14). Disruption of *tbc-2* decreased resistance to osmotic stress in *isp-1* worms (**E,F**; 450 mM NaCl, 550 mM NaCl), but resulted in an increased resistance to anoxia in *isp-1* mutants (**G**; 48 hours). Control data on wild-type and *tbc-2* mutants is repeated from Figure 2 for comparison. A minimum of three biological replicates were performed. Statistical significance was assessed using the log-rank test in panels A and D, a repeated measures ANOVA in panels B and C, and a two-way ANOVA with Šidák’s multiple comparison test in panels E-G. p values indicate significance of difference between *isp-1* (red line) and *isp-1;tbc-2* (purple line). Error bars indicate SEM. *p<0.05, ***p<0.001.

### Disruption of TBC-2 does not decrease *isp-1* lifespan and resistance to oxidative stress in a DAF-16 null background

Our previous work suggests that the effect of TBC-2 disruption on stress resistance in wild-type and *daf-2* worms is primarily dependent on DAF-16, while its effect on wild-type and *daf-2* lifespan is also mediated by TBC-2’s impact on DAF-16-independent pathways (Traa et al., 2022). To gain insight into the extent to which deletion of *tbc-2* decreases oxidative stress resistance and lifespan through DAF-16-dependent pathways, we examined the effect of disrupting *tbc-2* in *isp-1* worms lacking DAF-16 *(isp-1;daf-16* mutants). We found that *isp-1;daf-16* mutants have markedly decreased resistance to oxidative stress (4 mM paraquat) compared to *isp-1* worms and that this deficit was not significantly exacerbated by disruption of *tbc-2* (**Fig. 4A,B**). Similarly, *isp-1;daf-16* worms have a greatly reduced lifespan compared to *isp-1* worms and their lifespan is not further decreased by loss of *tbc-2* (**Fig. 4C,D**). In both cases, however, there was a trend towards decrease in the absence of *tbc-2*.

**Figure 4.**
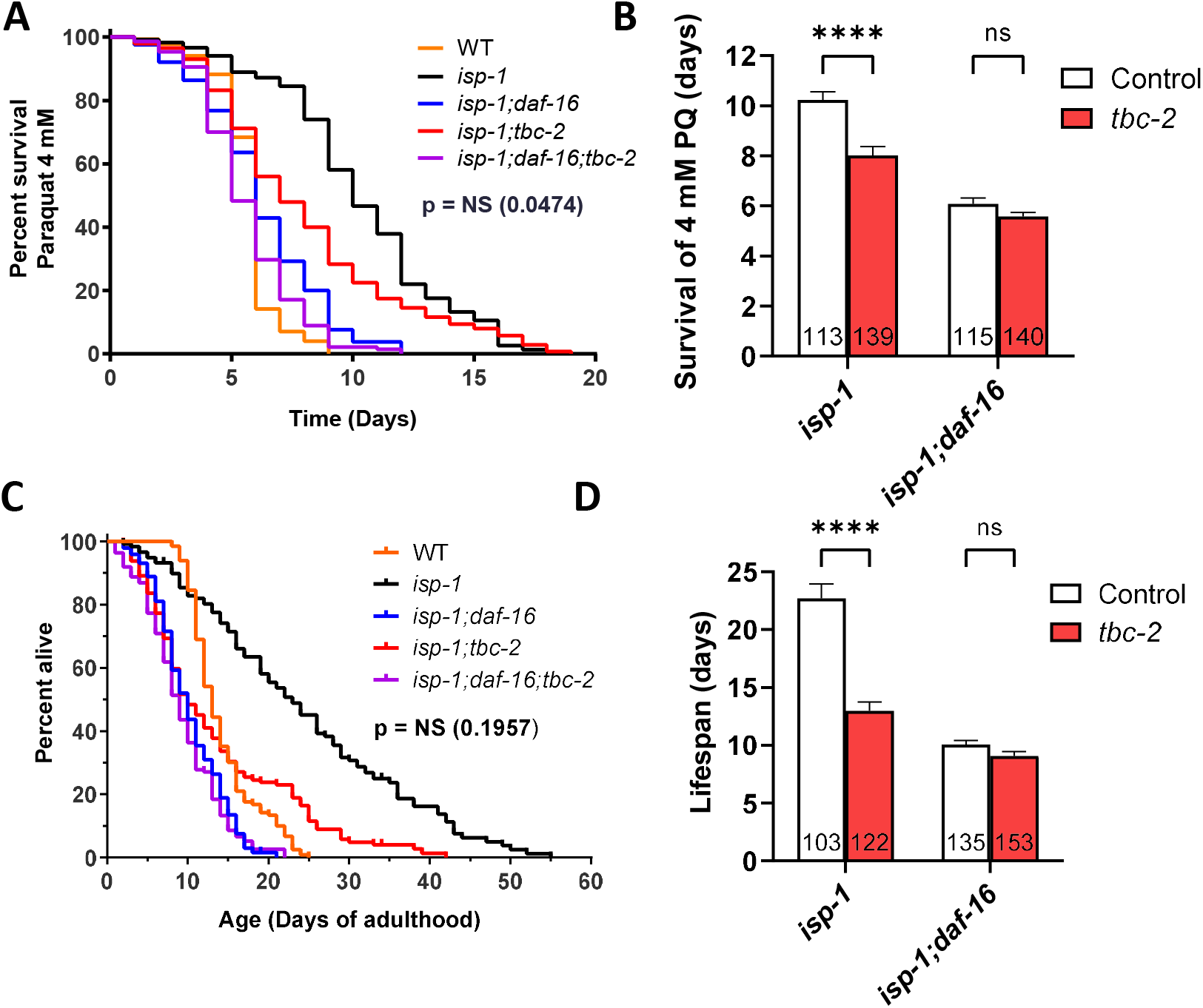
Loss of TBC-2 does not exacerbate the effects of disrupting DAF-16 on oxidative stress resistance or lifespan in *isp-1* mutants. To determine the role of DAF-16 in *tbc-2’s* detrimental impact on oxidative stress resistance and longevity, *tbc-2* was disrupted in *isp-1* worms with a deletion in *daf-16*. In *isp-1;daf-16* worms, deletion of *tbc-2* did not significantly decrease resistance to oxidative stress when exposed to 4 mM paraquat (**A,B**). Similarly, disruption of *tbc-2* did not significantly decrease *isp-1;daf-16* lifespan (**C,D**). In both cases, there was a trend towards decrease with the loss of *tbc-2*.Four biological replicates were performed. Statistical significance was assessed using a log-rank test in panels A and C, or a two-way ANOVA with Šidák’s multiple comparison test in panels B and D. In panels A and C, p value indicates difference between *isp-1;daf-16* and *isp-1;daf-16;tbc-2* and was corrected for multiple comparisons. Error bars indicate SEM. NS = not significant, ****p<0.0001.

### Disruption of TBC-2 does not prevent upregulation of DAF-16 target genes in long-lived mitochondrial mutants

In *daf-2* mutants, the upregulation of DAF-16 target genes is inhibited by the loss of *tbc-2* (Meras et al., 2022). Similarly, TBC-2 is required for the full upregulation of DAF-16 target genes in wild-type worms exposed to exogenous stressors (Traa et al., 2022). Since the long-lived mitochondrial mutants also exhibit upregulation of DAF-16 target genes (Senchuk et al., 2018), we wondered whether TBC-2 is required for the increased expression of DAF-16 targets in these mutants and if the prevention of this gene upregulation might be contributing to the decrease in stress resistance and lifespan resulting from *tbc-2* deletion. Accordingly, we used quantitative RT-PCR to compare the expression of DAF-16 target genes between long-lived mitochondrial mutants in a wild-type and *tbc-2* mutant background. For this purpose, we chose six DAF-16 target genes (*sod-3*, *dod-3*, *mtl-1*, *sodh-1*, *ftn-1*, *icl-1*) that were among the top fifty DAF-16 upregulated genes identified by a meta-analysis of DAF-16 gene expression studies (Tepper et al., 2013), and which we have previously used to examine DAF-16 activity (Campos et al., 2021; Meras et al., 2022; Senchuk et al., 2018; Traa et al., 2022).

Consistent with our past results (Senchuk et al., 2018), we found that all six DAF-16 target genes are upregulated in both *nuo-6* and *isp-1* mutants (**Fig. 5**). Disruption of *tbc-2* did not result in a strong or consistent effect on the expression of DAF-16 target genes in the long-lived mitochondrial mutants. There was one example in which loss of *tbc-2* significantly decreased DAF-16 target gene expression (*ftn-1* levels in *isp-1* mutants) and two examples in which the loss of *tbc-2* resulted in a significant increase in the expression of DAF-16 target genes (*dod-3* in *isp-1* worms and *sodh-1* in *nuo-6* mutants). Disruption of *tbc-2* did not significantly alter the expression of most of the DAF-16 target genes examined in wild-type, *nuo-6* or *isp-1* worms (**Fig. 5**). Combined, this suggests that disruption of *tbc-2* does not markedly inhibit the ability of *nuo-6* and *isp-1* worms to upregulate DAF-16 target genes. However, it is possible that the upregulation of other DAF-16 target genes in *nuo-6* and *isp-1* worms is affected by *tbc-2* disruption.

**Figure 5.**
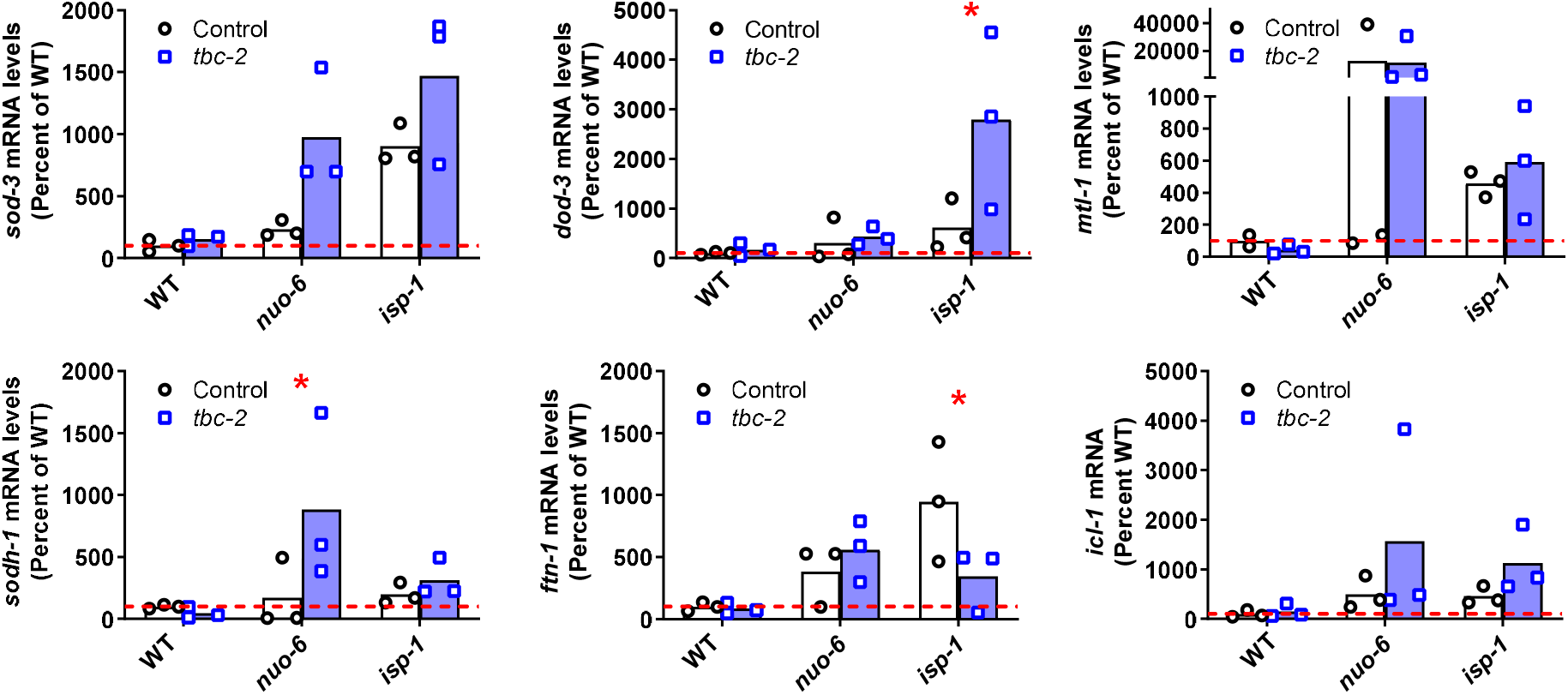
Disruption of TBC-2 does not prevent upregulation of DAF-16 target genes in long-lived mitochondrial mutants. Quantitative RT-PCR was used to quantify the effect of *tbc-2* deletion on the upregulation of DAF-16 target genes (*sod-3, dod-3, mtl-1, sodh-1, ftn-1,* and *icl-1)* in the long-lived mitochondrial mutants, *nuo-6* and *isp-1*. All of the DAF-16 target genes were upregulated in *nuo-6* and *isp-1* mutants. In all but one instance, the disruption of *tbc-2* did not prevent the upregulation of DAF-16 target genes in the long-lived mitochondrial mutants. This suggests that TBC-2 is not required for the upregulation of DAF-16 target genes in the long-lived mitochondrial mutants. Three biological replicates were performed. Statistical significance was assessed using a two-way ANOVA with Šidák’s multiple comparison test. Error bars indicates SEM. *p<0.05, **p<0.01, ***p<0.001.

## Discussion

In this work, we find that the loss of TBC-2 decreases both stress resistance and lifespan in *nuo-6* mutants and *isp-1* mutants. We previously showed that disruption of *tbc-2* also decreases stress resistance and lifespan in long-lived *daf-2* mutants (Traa et al., 2022). Interestingly, the pattern of decreased stress resistance resulting from deletion of *tbc-2* differs between these three strains (**Table S2**). While disruption of *tbc-2* decreases resistance to oxidative stress, heat stress, bacterial pathogen stress, osmotic stress and anoxia in *daf-2* mutants, resistance to heat stress, bacterial pathogen stress and anoxic stress is not decreased in either of the long-lived mitochondrial mutants when *tbc-2* is deleted. As our previous work suggests that the effect of *tbc-2* deletion on resistance to stress is primarily mediated by DAF-16 (Traa et al., 2022), it is plausible that stress resistance is more affected in *daf-2* mutants than *nuo-6* or *isp-1* mutants because the enhanced stress resistance of *daf-2* worms is entirely dependent on DAF-16 (Dues et al., 2019), while the long-lived mitochondrial mutants also rely on other pathways such as the mitochondrial unfolded protein response (Wu et al., 2018), the p38-mediated innate immune signaling pathway (Campos et al., 2021) and the mitochondrial thioredoxin system (Harris-Gauthier et al., 2022).

The only type of stress resistance that is significantly affected in all three strains is resistance to chronic oxidative stress. While this result is consistent with the possibility that disruption of *tbc-2* is decreasing lifespan by diminishing resistance to oxidative stress, the deletion of *tbc-2* also decreases lifespan in wild-type worms without affecting their ability to survive chronic oxidative stress (Traa et al., 2022). In addition, there are multiple examples in which resistance to oxidative stress can be experimentally dissociated from longevity (Shields et al., 2021; Van Raamsdonk and Hekimi, 2009; Van Raamsdonk et al., 2010).

To determine the extent to which deletion of *tbc-2* is decreasing oxidative stress resistance and lifespan through its effect on DAF-16, we examined the effect of disrupting *tbc-2* when DAF-16 is absent. We reasoned that if the *tbc-2* deletion is acting entirely through DAF-16 then disruption of *tbc-2* should not further decrease stress resistance or lifespan when DAF-16 is absent. We found that disruption of *tbc-2* resulted in a trend towards decreased resistance to chronic oxidative stress and lifespan in *isp-1;daf-16* mutants, which did not reach significance. This result is consistent with the effects of *tbc-2* deletion being mediated by DAF-16. In contrast, we found that disruption of *tbc-2* did not prevent the upregulation of DAF-16 target genes in the long-lived mitochondrial mutants, which is consistent with the effects of *tbc-2* deletion being mediated by DAF-16-independent pathways. These differing conclusions may stem from the *daf-16* deletion having such a severe impact on *isp-1* stress resistance and longevity that the effects of *tbc-2* disruption on DAF-16-independent pathways are being masked. This possibility is supported by the fact that we did observe a trend towards decrease in *isp-1;daf-16;tbc-2* mutants compared to *isp-1;daf-16* mutants.

In contrast to the long-lived mitochondrial mutants, TBC-2 is required for the full upregulation of DAF-16 target genes in *daf-2* worms (Meras et al., 2022) and also important for the nuclear localization of DAF-16 in *daf-2* mutants (Traa et al., 2022). In these worms, the disruption of TBC-2 appears to be affecting stress resistance primarily through DAF-16 but affecting lifespan through DAF-16 and DAF-16-independent pathways (Traa et al., 2022). These differences likely stem from *daf-2* mutants being more dependent on DAF-16 for their stress resistance and longevity than the long-lived mitochondrial mutants, which rely more on DAF-16-independent pathways compared to *daf-2* worms. These DAF-16-independent pathways may include signaling through the nuclear hormone receptor NHR-49, epidermal growth factor signaling and notch signaling, all of which have been shown to be present on endosomes and influence longevity (Arantes-Oliveira et al., 2002; Curran and Ruvkun, 2007; Dresen et al., 2015; Fortini and Bilder, 2009; Iwasa et al., 2010; Khan et al., 2013; Walter et al., 2011; Wang et al., 2002; Watterson et al., 2022; Yu and Driscoll, 2011; Zheng et al., 2013). In the future, it will be important to assess the contribution of these and other signaling pathways that are present on endosomes in the long-lived mitochondrial mutants.

Two other endosome related proteins were recently shown to be required for the longevity of *isp-1* worms. After using proteomics to identify differentially expressed proteins in long-lived *clk-1* and *isp-1* mutants, a targeted RNAi screen was performed to assess the contribution of these proteins to their longevity. It was found that RNAi knockdown or genetic deletion of *mon-2* decreases *isp-1* lifespan (Jung et al., 2021). MON-2 regulates trafficking between endosomes and the Golgi apparatus (Mahajan et al., 2013). Similarly, TBC-3 is also involved in transport between endosomes and Golgi (Kanamori et al., 2008) and is required for *isp-1* lifespan (Jung et al., 2021). While MON-2 appears to be promoting longevity by upregulating autophagy, it is unclear how elevated endosomal MON-2 increases autophagy (Artan et al., 2022; Jung et al., 2021). Future studies will be needed to determine whether MON-2 and TBC-3 are required for endosomal signaling pathways, similar to TBC-2, or whether these endosome-related proteins are all acting through independent mechanisms.

## Conclusions

We find that disruption of the endosomal trafficking protein TBC-2 decreases oxidative stress resistance and lifespan in the long-lived mitochondrial mutants *nuo-6* and *isp-1*. Interestingly, the loss of TBC-2 does not prevent the upregulation of DAF-16 target genes in these mutants suggesting that the disruption of *tbc-2* may be affecting lifespan and resistance to stress in these worms by affecting other signaling pathways localized to endosomes. This work demonstrates an important role of endosomal signaling in stress resistance and lifespan.

## Materials and Methods

### Strains

N2 (WT)

QR15 *tbc-2(tm2241) II*

JVR171 *isp-1(qm150) IV*

MQ1333 *nuo-6(qm200) I*

JVR555 *isp-1(qm150) IV;tbc-2(tm2241)*

JVR556 *nuo-6(qm200) I;tbc-2(tm2241)*

JVR502 *daf-16(mu86) I;isp-1(qm150) IV*

JVR596 *daf-16(mu86) I;tbc-2(tm2241) II*

JVR619 *daf-16(mu86) I;tbc-2(tm2241) II;isp-1(qm150) IV*

Worms were maintained at 20°C on NGM plates seeded with OP50 bacteria.

### Heat Stress Assay

Resistance to heat stress was measured by transferring 25 pre-fertile young adult worms to freshly seeded NGM plates, incubating at 37°C and monitoring survival every 2 hours for a total of 10 hours.

### Chronic Oxidative Stress Assay

Resistance to chronic oxidative stress was measured by transferring 30 pre-fertile young adult worms to 60 mm NGM plates containing 4 mM paraquat (methyl viologen, Sigma Catalog No. 856177) and 100 μM FUdR and seeded with concentrated OP50 bacteria (see (Senchuk et al., 2017) for detailed protocol). Worms were kept at 20°C and survival was monitored daily until all worms died.

### Acute Oxidative Stress Assay

Resistance to acute oxidative stress was measured by transferring 30 pre-fertile young adult worms to 300 μM juglone plates seeded with OP50 bacteria and monitoring survival at 20°C every 2 hours for a total of 8 hours. Plates were made fresh on the day of the assay as juglone toxicity diminishes rapidly with time (see http://wbg.wormbook.org/2016/07/14/measuring-sensitivity-to-oxidative-stress-the-superoxide-generator-juglone-rapidly-loses-potency-with-time/).

### Osmotic Stress Assay

To measure resistance to osmotic stress, 25 pre-fertile young adult worms were transferred to 450 or 550 mM NaCl plates seeded with OP50 bacteria and survival was scored after 24 hours at 20°C.

### Anoxia Assay

To measure resistance to anoxic stress, 45 pre-fertile young adult worms were transferred to NGM plates freshly seeded with OP50 bacteria and sealed in low-oxygen Becton-Dickinson Bio-Bag Type A Environmental Chambers. After 48 hours at 20°C, worms were allowed to recover outside of the chamber for 24 hours before survival was measured.

### Bacterial Pathogen Stress Assay

Resistance to bacterial pathogen stress was performed as previously described (Campos et al., 2021). 45 L4 worms were transferred to 100 mg/L FUdR plates seeded with OP50 bacteria and were grown at 20°C until day 3 of adulthood. Day 3 adult worms were then transferred from these plates to 20 mg/L FUdR plates seeded with *Pseudomonas aeruginosa* (PA14). Worms were kept at 20°C and survival was monitored daily until all worms died.

### Lifespan Assay

To measure lifespan, 50 pre-fertile young adult worms were transferred to 25 μM FUdR plates seeded with OP50 bacteria and kept at 20°C. Low concentrations of FUdR limit internal hatching and inhibit development of progeny without affecting longevity in wild-type worms (Van Raamsdonk and Hekimi, 2011). Survival was monitored daily until all worms died.

### Quantification of mRNA levels by quantitative RT-PCR

Quantitative RT-PCR was performed by collecting worms in M9 buffer. RNA was extracted using Trizol and converted to cDNA using a high-capacity cDNA Reverse Transcription kit (Applied Biosystems 4368814). Quantitative PCR was performed using a PowerUp SYBR Green Master Mix (Applied Biosystems A25742) in a MicroAmp Optical 96-well reaction plate (Applied Biosystems N8010560) and a Viia 7 applied biosystems qPCR machine. mRNA levels were calculated as the copy number of the gene of interest relative to the copy number of the endogenous control, *act-3,* and expressed as a percentage of WT. Primer sequences for each target gene are as follows:

*sod-3* (AAAGGAGCTGATGGACACTATTAAGC, AAGTTATCCAGGGAACCGAAGTC),
*dod-3* (AAGTGCTCCGATTGTTACGC, ACATGAACACCGGCTCATTC),
*mtl-1* (ATGGCTTGCAAGTGTGACTG, GCTTCTGCTCTGCACAATGA),
*sodh-1* (GAAGGAGCTGGAAGTGTTGTTC, CTCCACGTATAGTGAGGTACTCCTG),
*ftn-1* (GAGTGGGGAACTGTCCTTGA, CGAATGTACCTGCTCTTCCA),
*icl-1* (TGTGAAGCCGAGGACTACCT, TCTCCGATCCAAGCTGATCT),
*act-3* (TGCGACATTGATATCCGTAAGG, GGTGGTTCCTCCGGAAAGAA).

### Statistical Analysis

At least three biological replicates were completed in each experiment. For lifespan and stress assays, the experimenter was blinded to the genotype to ensure unbiased results. Worms were excluded from lifespan and stress assays if they crawled off the agar and died on the side of the plate, had internal hatching of progeny or expulsion of internal organs. Statistical significance of differences between genotypes was determined using a two-way ANOVA with Šidák’s multiple comparison test, a repeated measures ANOVA, or a log-rank test using Graphpad Prism.

## Acknowledgments

We would like to thank Christian Rocheleau for discussions about this work and for reviewing the manuscript prior to submission. Some strains were provided by the CGC, which is funded by NIH Office of Research Infrastructure Programs (P30 OD010440). We would also like to acknowledge the *C. elegans* knockout consortium and the National Bioresource Project of Japan for providing strains used in this research.

## Author Contributions

Conceptualization: JVR. Methodology: AT, HS, AA, ZR, BK, JVR. Investigation: AT, HS, AA, ZR, BK. Analysis: AT, HS, AA, ZR, BK, JVR. Visualization AT, HS, AA, ZR, BK, JVR. Writing – original draft: AT, JVR. Writing – review and editing: AT, HS, AA, ZR, BK, JVR. Supervision: JVR.

## Materials & Correspondence

Correspondence and material requests should be addressed to Jeremy Van Raamsdonk.

---

**Table S2.** Effect of TBC-2 disruption on stress resistance, gene expression and lifespan.

|  | **Chronic**<br>**Oxidative**<br>**Stress** | **Acute**<br>**Oxidative**<br>**Stress** | **Heat**<br>**Stress** | **Bacterial**<br>**Pathogen**<br>**Stress** | **Osmotic**<br>**Stress** | **Anoxia** | **Expression of**<br>**DAF-16 Target**<br>**Genes** | **Lifespan** |
| --- | --- | --- | --- | --- | --- | --- | --- | --- |
| **WT** | No effect | Decreased | Decreased | Decreased | No effect | Decreased | No effect | Decreased |
| ***nuo-6*** | Decreased | Decreased | No effect | No effect | No effect | No effect | No effect | Decreased |
| ***isp-1*** | Decreased | No effect | No effect | Increased | Decreased | Increased | No effect | Decreased |
| ***daf-2*** | Decreased | No effect | Decreased | Decreased | Decreased | Decreased | Decreased | Decreased |

